# Synaptic polarity and sign-balance prediction using gene expression data in the *Caenorhabditis elegans* chemical synapse neuronal connectome network

**DOI:** 10.1101/2020.05.22.110312

**Authors:** Bánk G. Fenyves, Gábor S. Szilágyi, Zsolt Vassy, Csaba Sőti, Péter Csermely

## Abstract

Graph theoretical analyses of nervous systems usually omit the aspect of connection polarity, due to data insufficiency. The chemical synapse network of *Caenorhabditis elegans* is a well-reconstructed directed network, but the signs of its connections are yet to be elucidated. Here, we present the gene expression-based sign prediction of the *C. elegans* connectome, incorporating presynaptic neurotransmitter and postsynaptic receptor gene expression data (3,638 connections and 20,589 synapses total). We made successful predictions for more than two-thirds of all chemical synapses and determined a ratio of excitatory-inhibitory (E:I) interneuronal ionotropic chemical connections close to 4:1 which was found similar to that observed in many real-world networks. Our open source tool (http://EleganSign.linkgroup.hu) is simple but efficient in predicting polarities by integrating neuronal connectome and gene expression data.

**Author Summary:** The fundamental way neurons communicate is by activating or inhibiting each other via synapses. The balance between the two is crucial for the optimal functioning of a nervous system. However, whole-brain synaptic polarity information is unavailable for any species and experimental validation is challenging. The roundworm Caenorhabditis elegans possesses a fully mapped connectome with a comprehensive gene expression profile of its 302 neurons. Based on the consideration that the polarity of a synapse must be determined by the neurotransmitter(s) expressed in the presynaptic neuron and the receptors expressed in the postsynaptic neuron, we conceptualized and created a tool that predicts synaptic polarities based on connectivity and gene expression information. We were able to show for the first time that the ratio of excitatory and inhibitory synapses in *C. elegans* is around 4 to 1 which is in line with the balance observed in many natural systems. Our method opens a way to include spatial and temporal dynamics of synaptic polarity that would add a new dimension of plasticity in the excitatory:inhibitory balance. Our tool is freely available to be used on any network accompanied by any expression atlas.

## Introduction

**C**hemical synapses of a neuronal network are both directed and signed, since a neuron is able to excite or inhibit another neuron. The nervous system of the nematode *Caenorhabditis elegans* has been fully mapped and reconstructed [1–3]. However, except for a few connections there is no comprehensive chemical synapse polarity data available [1]. While the direction of a synaptic connection can be inferred from its structure, the experimental determination of its polarity requires delicate electrophysiological methods (e.g. patch-clamping) which have a limited system-level use. Instead, *in silico* approaches using reverse engineering have efficiently predicted synaptic signs for subnetworks of the *C. elegans* connectome [4–6].

Many synaptic sign prediction models have relied on the widely accepted assumption that the polarity of a chemical synapse is solely determined by the type of neurotransmitter released by the presynaptic neuron [4,5]. Therefore, in *C. elegans* excitatory glutamatergic and cholinergic, as well as inhibitory γ-aminobutyric acid (GABA)-ergic ionotropic chemical connections have been modeled. However, with this approach, approximately 95% of the connections turned to be excitatory [7,8]. This would possibly result in an unbalanced, over-excited network, as has been shown by previous publications [9–11]. Moreover, there is evidence of unconventional postsynaptic effects of neurotransmitters, such as cholinergic inhibition [12,13] or glutamatergic inhibition [14–16], meaning that a neuron can simultaneously excite and inhibit its postsynaptic partners with the same neurotransmitter due to variable neurotransmitter receptor expression on the postsynaptic neuron membrane. For example, the cholinergic AIY interneuron can activate RIB neurons and inhibit AIZ neurons in an acetylcholine-mediated fashion [13].

We aimed to predict synaptic polarities of ionotropic chemical connections of the *C. elegans* connectome relying on presynaptic and postsynaptic gene expression data. In this study of the *C. elegans* nervous system, we successfully predicted the polarity of more than 70% of ionotropic chemical synapses and predicted a sign-balance of excitatory:inhibitory connections close to 4:1 that was observed and functionally stable in many real-world circumstances. Presenting the first connectome-scale work, we show that the concept of gene expression-based polarity prediction can efficiently be applied to demonstrate a natural balance in a nervous system.

## Results

### Creating a prediction tool of the synaptic polarities of the *C. elegans* connectome

Our primary goal was to infer synaptic polarity from combining connectivity and gene expression data. We created a simple, yet powe algorithmic database (S1 Data) that takes as input connectome and gene expression data to predict synaptic polarity of ionotropic chemical connections. We used the *C. elegans* WormWiring connectome data primarily in the form of a weighted edge list representing 20,589 chemical synapses in 3,638 connections. Our prediction tool is available here: http://EleganSign.linkgroup.hu.

### Update of the previous neurotransmitter expression tables

We updated the *C. elegans* neuronal neurotransmitter tables previously published [17–20] with recent evidence [8,21] (Methods). After this update, 256 neurons had a single neurotransmitter expressed, while 12 neurons had double neurotransmitter expression (Fig 1A). There were 34 neurons which did not express any of the three neurotransmitters investigated.

**Fig 1.**
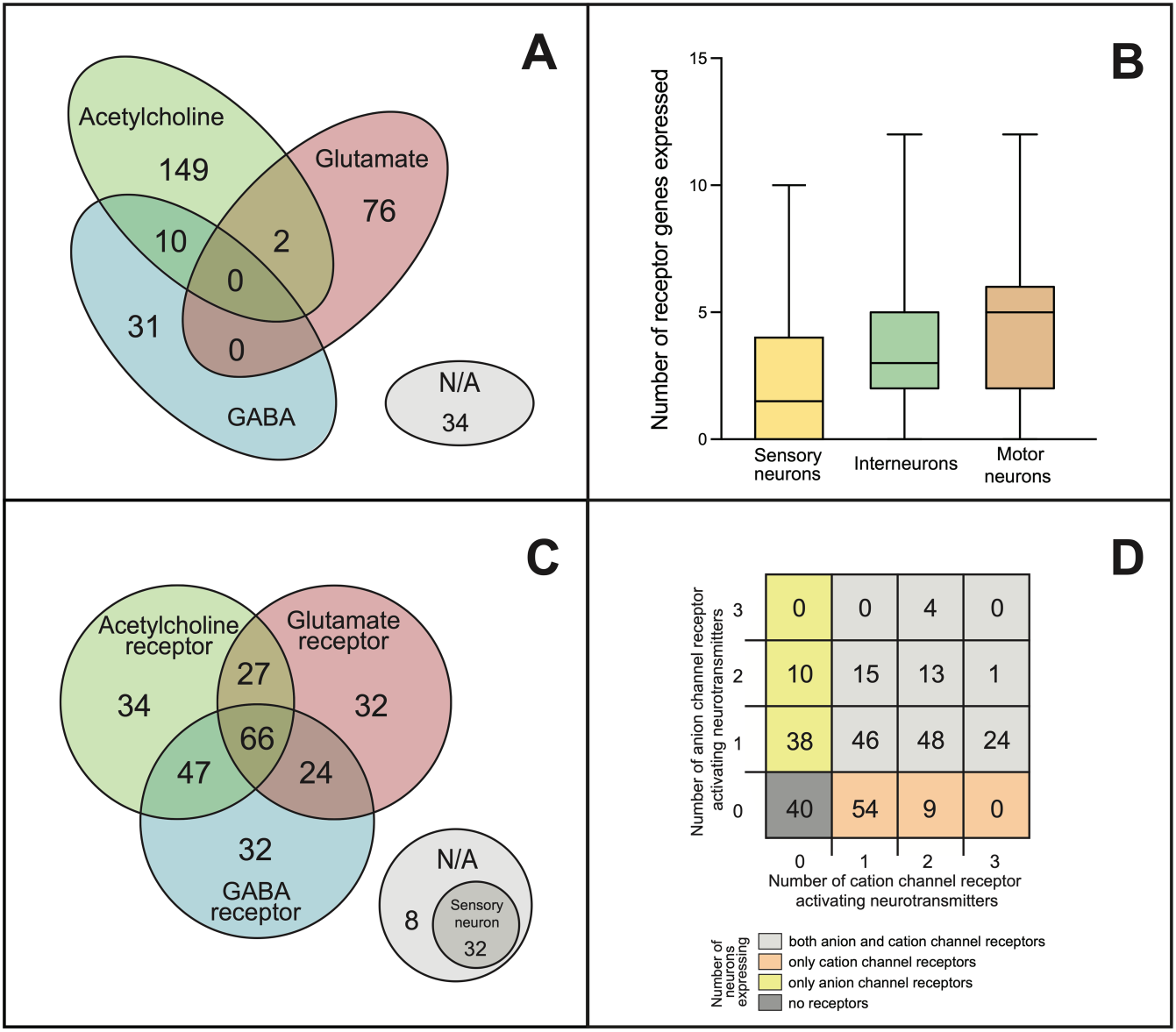
Neurotransmitter and receptor expression patterns of *C. elegans* neurons. Expression data of the three major synaptic neurotransmitters and their receptors of *C. elegans* were collected from multiple datasets and were manually curated (see Methods). **(A)** Distribution of neurons according to their neurotransmitter expression: glutamate (red), acetylcholine (green), GABA (blue) or none (grey). **(B)** Distribution of neurons based on their neurotransmitter receptor gene expression (colors are the same as in panel **a**). **(C)** Number of receptor genes expressed by neurons, grouped by neuron modality. **(D)** Distribution of neurons according to the number of neurotransmitters for which anion and/or cation channel receptor genes are expressed.

### Extraction of gene expression data

In parallel, we extracted gene expression data from Wormatlas [22], Wormbase (www.wormbase.org), and a recent RNA-sequencing dataset [21] which we curated manually to assign ionotropic receptor expression pattern to each neuron (see Methods). To do this we first sorted the previously identified 62 ionotropic receptor genes into six functional classes based on their suggested neurotransmitter ligand (glutamate, acetylcholine or GABA) and putative ion channel type (cationic or anionic, i.e. excitatory or inhibitory), as shown in Table 1. We found evidence for expression of 42 out of the 62 receptor genes in the *C. elegans* nervous system (genes marked bold in Table 1; also S1 Data). We also found an increasing average number of receptor genes in sensory, inter- and motor neurons, respectively (Fig 1B).

**Table 1.** Neurotransmitter receptor genes.

|  | Glutamate | Acetylcholine | GABA |
| --- | --- | --- | --- |
| cation channel<br>receptor gene | *glr-1*<br>*glr-2*<br>*glr-3*<br>*glr-4*<br>*glr-5*<br>*glr-6*<br>*glr-7*<br>*glr-8*<br>*nmr-1*<br>*nmr-2* | *acr-1* *acr-16*<br>*acr-2* *acr-17*<br>*acr-3* *acr-18*<br>*acr-4* *acr-19*<br>*acr-5* *acr-20*<br>*acr-6* *acr-21*<br>*acr-7* *acr-23*<br>*acr-8* *acr-25*<br>*acr-9* *deg-3*<br>*acr-10* *des-2*<br>*acr-11* *lev-8*<br>*acr-12* *unc-29*<br>*acr-13* *unc-38*<br>*acr-14* *unc-63*<br>*acr-15* | *exp-1*<br>*lgc-35* |
| **anion channel**<br>**receptor gene** | *glc-1* | *acc-1* | ***gab-1*** |
|  | *glc-2* | *acc-2* | ***ggr-1*** |
|  | ***glc-3*** | *acc-3* | ***ggr-2*** |
|  | *glc-4* | ***lgc-46*** | *ggr-3* |
|  | ***avr-14*** | ***lgc-47*** | ***lgc-36*** |
|  | ***avr-15*** | *lgc-48* | ***lgc-37*** |
|  |  | *lgc-49* | ***lgc-38*** |
|  |  |  | *unc-49* |

The *Caenorhabditis elegans* genome contains 62 ionotropic postsynaptic receptor genes for glutamate, acetylcholine, and GABA. In this table genes are grouped according to their neurotransmitter ligand and whether forming cationic (+) or anionic (-) ion channels ([22] and Table 1 in Supporting Information). In *C. elegans* "unconventional signaling", namely, glutamate-mediated inhibition, cholinergic inhibition and GABA-ergic excitation, is facilitated by 6, 7, and 2 receptor genes, respectively. In the gene expression database used in this work, expression in at least one neuron was found in the case of 42 genes (marked **bold**), while for 20 genes no neuronal expression was found.

Next, for all the 302 neurons of the *C. elegans* connectome we determined which receptor classes were expressed. 164 neurons had an overlapping expression of receptors for two or three different neurotransmitters (Fig 1C). The distribution of neurons according to their expression of cationic and/or anionic glutamate, acetylcholine and/or GABA receptors suggested functional diversity due to the high number of neurons expressing both excitatory and inhibitory receptors (Fig 1D). Surprisingly, 85 neurons expressed both excitatory and inhibitory receptor genes for the same neurotransmitter (S1 Data). Forty out of 302 neurons showed no receptor expression, out of which 32 neurons were primarily sensory neurons (S1 Data). The average number of receptor genes expressed was 3.6 per neuron (Table 2 in S1 Supporting Information).

**Table 2.** Proportions of negative edges in signed networks.

| Network | Proportion of negative edges | Reference |
| --- | --- | --- |
| ***C. elegans* chemical synapse network** | **20%** | **this work** |
| **Rat hippocampus (*in vivo*)** | **5-30%** | [33] |
| **Rat excitatory neocortical neurons** | **20%** | [34] |
| **Cerebral cortex (*in vivo*; GAD expression)** | **10-20%** | [15] |
| **Cerebral cortex (*in vivo*; GABA neurons)** | **20-25%** | [15] |
| **Optimal network for synchronized bursting activity (*in silico*)** | **10-20%** | [6] |
| **Primary visual cortex (V1; *in silico*)** | **25%** | [44] |
| **Neuronal network (*ex vivo*) and neuronal network model (*in silico*)** | **20%** | [16] |
| Wikipedia (social network) | 21% | [35] |
| Epinions (social network) | 15% | [35] |
| Slashdot (social network) | 23% | [35] |
| University freshman network (social) | 12-14% | [36] |

### Neurotransmitter and receptor gene expression-based polarity prediction

After assigning neurotransmitter and receptor expression patterns to each neuron, we predicted synaptic polarities by looking for matches between the neurotransmitter expression of the presynaptic neuron and the receptor gene expression of the postsynaptic neuron (Fig 2A). This way, we labeled synapses as one of the following: excitatory, inhibitory, complex, or unpredicted (see Methods). “Excitatory” or “inhibitory” label were given when the neurotransmitter-matched postsynaptic receptor genes were only cation or anion channel related, respectively. A synapse was labeled as “complex” if data suggested both excitatory and inhibitory function. With this approach, we successfully predicted synaptic polarity for 73% of chemical synapses of the *C. elegans* connectome (Fig 2B). We couldn’t predict polarity for the remaining synapses due to missing neurotransmitter data or mismatch in neurotransmitter/receptor expression (Fig 2B). We predicted that 9,034 of the synapses are excitatory and 2,580 are inhibitory, while 3,431 synapses have complex function (Fig 3A; S1 Data). These findings suggest that the overall ratio of excitatory and inhibitory synapses (E:I ratio) in the *C. elegans* ionotropic chemical synapse network is close to 4:1 (Fig 3A, *NT*+*R method*).

**Fig 2.**
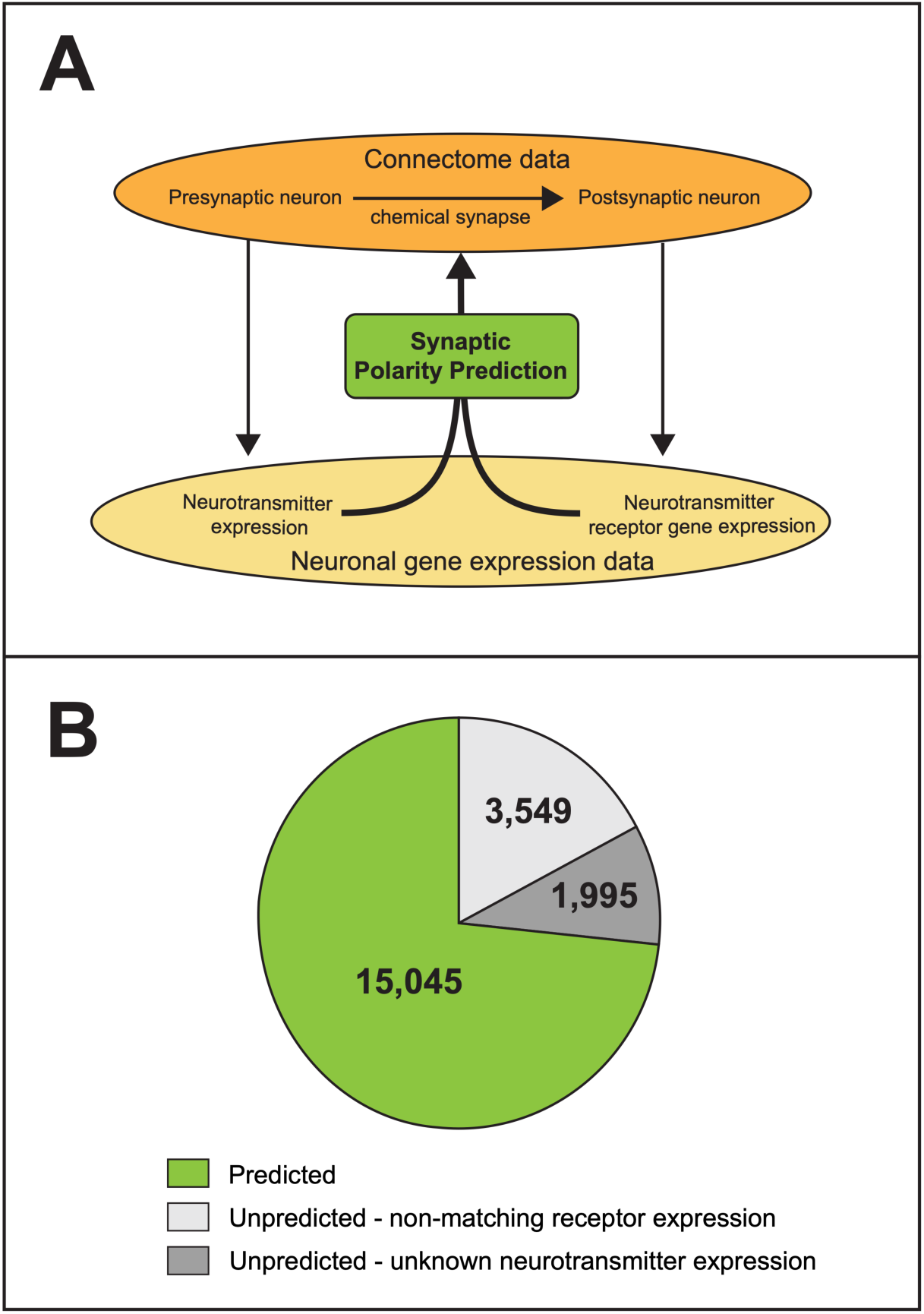
Prediction of synaptic polarities of the *C. elegans* connectome. **(A)** Prediction method. Connectome and gene expression data were manually curated (see Methods). Polarities of chemical synapses were predicted based on the neurotransmitter expression of presynaptic neurons and the matching receptor gene expression of the postsynaptic neurons. **(B)** Distribution of predicted and unpredicted synapses. We were able to predict the polarity of 73% of chemical synapses (green). The polarities of the rest of synapses were unpredicted due to unknown neurotransmitter expression of the presynaptic neurons (dark grey) or non-matching receptor gene expression of the postsynaptic neurons (light grey).

**Fig 3.**
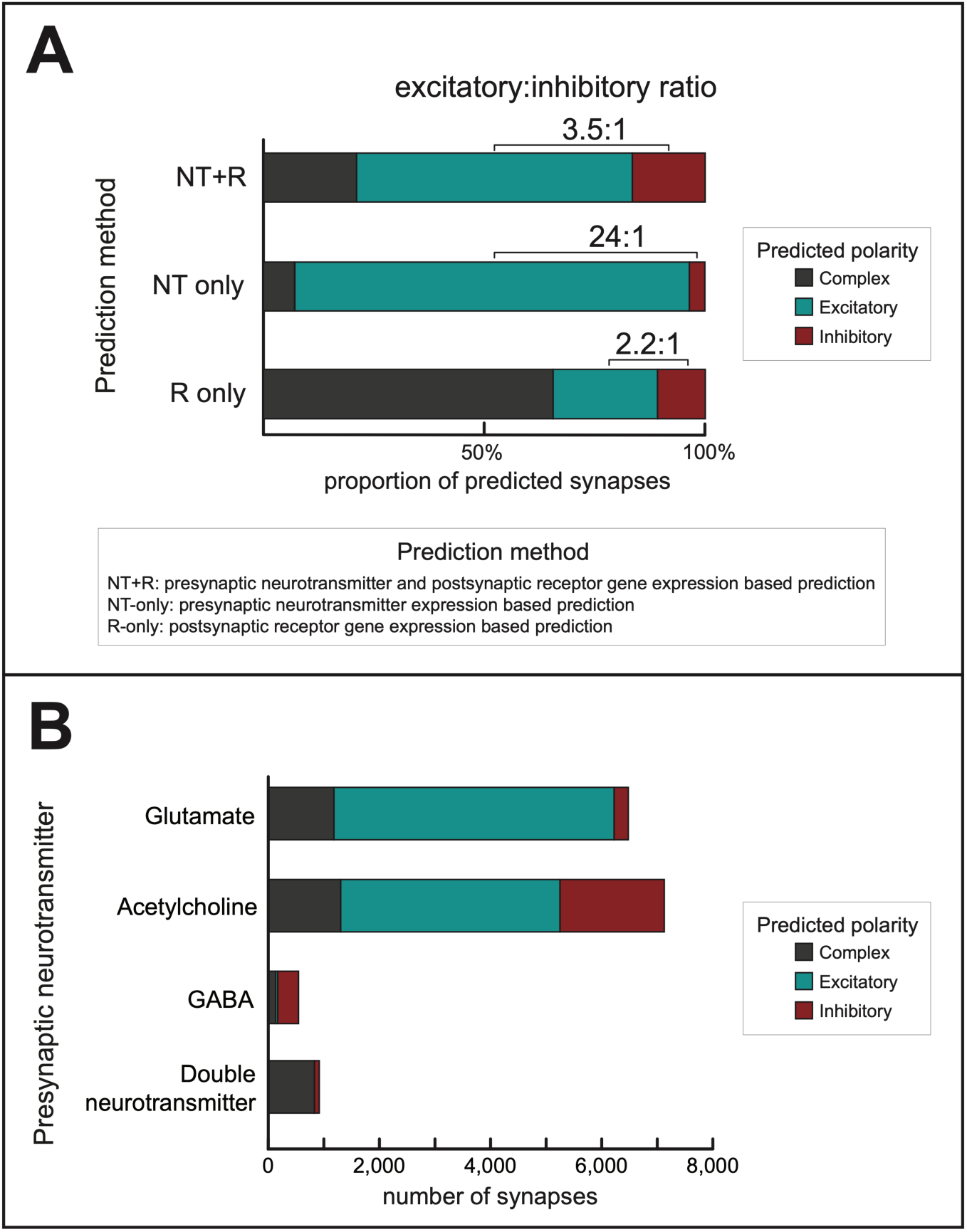
Predicted synaptic polarities. **(A)** Distributions of predicted polarities, using the method developed in this paper (*NT*+*R*) and two alternative methods as comparison (*NT-only* and *R-only*). Polarities were predicted by considering the neurotransmitter expression of the presynaptic neuron and/or the receptor gene expression of the postsynaptic neuron (see Methods). **(B)** Distributions of predicted synaptic polarities (using the *NT*+*R method*) according to the presynaptic neurotransmitter. Colors are the same as in panel **A**. Unpredicted synapses are not shown.

### Alternative polarity prediction methods

To put our results in context, we applied two alternative prediction methods for comparison (*NT-only* and *R-only*; see Methods). The *NT-only* method yielded a much higher E:I ratio (Fig 3A, *NT-only method*; S2 Data), which is in line with the dominance of purely glutamatergic or cholinergic (traditionally excitatory) neurons over GABA-ergic (traditionally inhibitory) neurons (Fig 1A). To explain the difference further, the *NT*+*R* method predicted that 30% of cholinergic and 5% of glutamatergic synapses were *inhibitory* (Fig 3B) which is a significant fraction of synapses otherwise predicted excitatory with the *NT-only* method. The other, *R-only* method yielded a markedly smaller E:I ratio, however predicted an excessive number of complex synapses (Fig 3A, *R-only method*, S3 Data). This is due to the fact that many neurons express both cation and anion channel receptor genes (Fig 1D).

### Feedback inhibition between neuron groups

Notably, in subsets of connections which connect neurons of different modalities of sensory neurons, motor neurons, interneurons and polymodal neurons, the E:I ratios varied between 1:10 (motor –> sensory) and 14:1 (inter –> motor). Importantly, we observed dominant inhibition in the motor –> sensory, motor –> inter, and inter –> sensory directions (Fig 4A), exhibiting inhibitory backwards signaling as discussed in the literature previously [23,24]. In addition, significant presence of inhibitory and complex connections was found in the locomotion circuit as well (Fig 4B).

**Fig 4.**
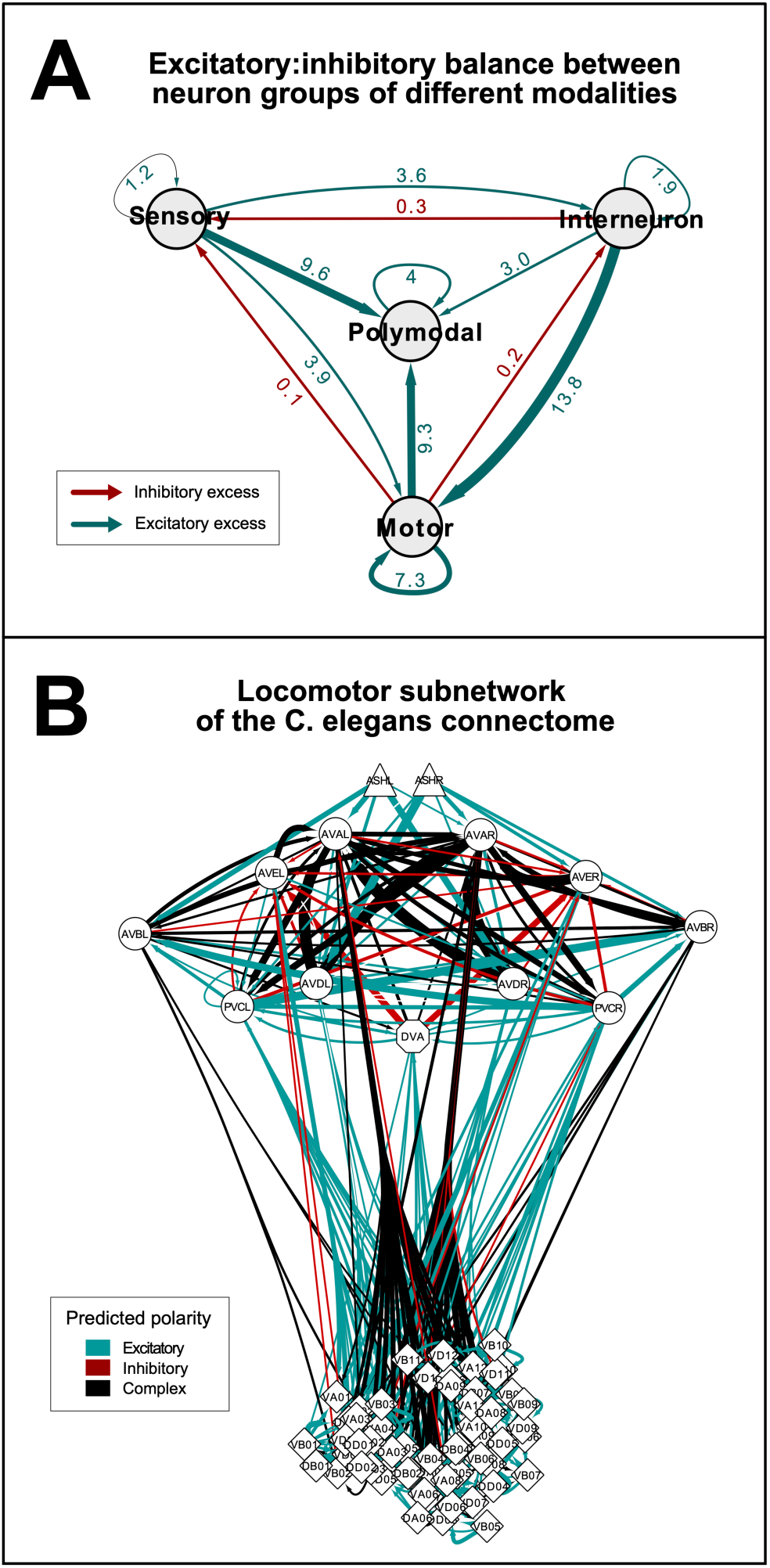
Excitatory:inhibitory balance of different neuron groups. **(A)** Excitatory:inhibitory balance between neuron groups of different modalities. Nodes represent groups of neurons by modality. Edges are weighted according to the excitatory:inhibitory (E:I) ratios (see numbers). Green and red colors represent excitatory (E:I>1) and inhibitory (E:I<1) excess in sign-balances, respectively. **(B)** Network representation of the locomotion subnetwork. Edges represent excitatory (blue), inhibitory (red), or complex (black) chemical connections. Edges are weighted according to synapse number. Shape of vertices (Δ,○,◇) represent the modality (sensory, inter, motor, respectively) of neurons.

### Network representations of the signed *C. elegans* connectome

Network representations of synaptic polarities in the *C. elegans* chemical synapse connectome using the EntOptLayout plugin of Cytoscape [25] are on Fig 5. Fig 5A shows that the modular structure of the *C. elegans* connectome visualized by the EntOptLayout method nicely captures the anatomical locations of the anterior, ventral and lateral ganglions, as well as the premotor interneurons of the worm. While the anterior and lateral ganglions show a large glutamate expression, this is much less characteristic to the ventral ganglion (Fig 5A). Fig 5B shows that the ventral ganglion has predominantly inhibitory connections, while connections in the other locations are predominantly excitatory if predicted by our *NT* + *R method*. Fig 5C demonstrates that the prediction of using neurotransmitters only (*NT-only method*) results in a large excitatory excess, which makes the system unstable. The anatomical locations of neurons expressing various neurotransmitters (Fig 5D) correspond well to the network representation shown on Fig 5A. Links between inhibitory function and anatomical structures have been shown in the human brain [26,27], but have not been demonstrated previously in the nematode.

**Fig 5.**
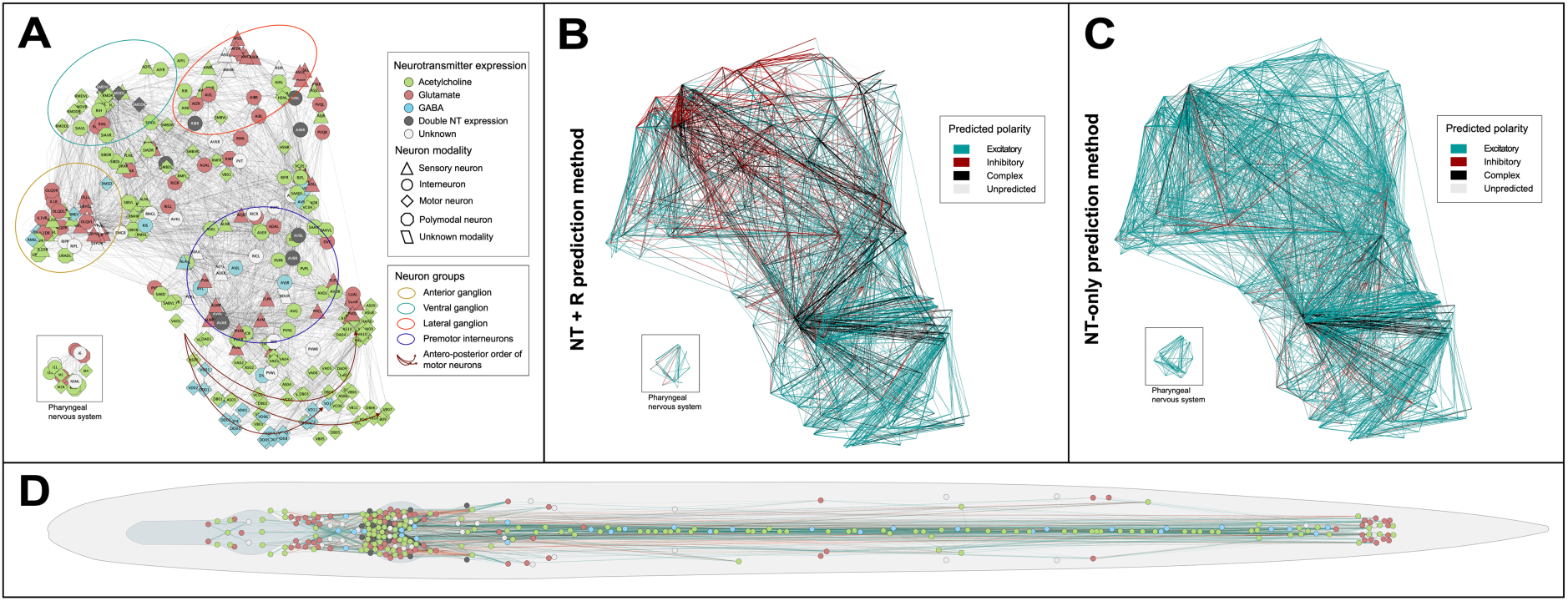
Network representations of the *C. elegans* chemical synapse network. **(A-C)** Network representations using the EntOpt layout plugin in Cytoscape[25]. **(A)** Color and shape of vertices represent neurotransmitter expression and modality of neurons, respectively (see inset for definitions). **(B)** Edges represent excitatory (blue), inhibitory (red), or complex (black) chemical connections predicted by the NT+R method (see Methods), weighted according to synapse numbers. **(C)** Colors of edges (see panel **B**) represent the polarities of chemical synapses predicted by the NT-only method. **(D)** Layout of vertices is representing the anatomic position of neurons. Node and edge colors are as in panels **A** and **B**, respectively.

### Validation of our predictions

To validate our results, we compared them with an *in silico* prediction work of the locomotion circuit of *C. elegans* [28] and confirmed that 70% of the synapses were predicted the same (Table 3 in S1 Supporting Information), albeit using a completely different concept. As a further validation step, we tested our predictions against experimental evidence and found that the majority of predicted polarities using our method were consistent with earlier findings in *C. elegans*: only a single interneuronal connection out of 12 showed opposing polarity (Table 4 in S1 Supporting Information).

To test the robustness of our *NT*+*R* prediction method, we applied the same rules to predict polarities of connections regardless of synapse number data, and after perturbations in the network like deletion of the 20 pharyngeal nervous system neurons or deletion of potentially variable (i.e. single-synapse) connections (S4 Data). Furthermore, we repeated our analyses in other published connectomes [2,3] of different sizes as well (Table 5 in S1 Supporting Information and S5 Data). In all five cases, the excitatory:inhibitory ratios were in the range of 3.1 to 4.1 (S1 Fig). When predicting not using the yet preprint-published RNA-seq expression dataset [21], this range was 4.1-4.6 (data not shown). All together, these findings suggest that the observed sign-balance is a remarkably robust property of the *C. elegans* neuronal network.

## Discussion

Nervous systems are not only directed but signed networks as well, since neurons either activate or inhibit other neurons [10]. The balance of excitatory and inhibitory connections (i.e. the sign-balance) is a fundamental feature of brain networks, clearly marked by the variety of disorders associated with its impairment [10,29,30]. However, direct evidence of single synapse polarity is rather sporadic even in simple species. In this work we predicted that the sign-balance in the *C. elegans* chemical synapse network is approximately 4:1 (excitatory-inhibitory, E:I). This is consistent with previous *in vitro* and *in vivo* studies of nervous systems [31–34], and also with observations of different social networks [35,36], as shown in Table 2. However, this ratio can only be predicted if not only the neurotransmitter expression of the presynaptic neuron but also the receptor gene expression of the postsynaptic neuron is taken into consideration. Its significance is due to fifteen receptor genes which are presumed to encode inhibitory glutamatergic/cholinergic or excitatory GABA-ergic ion channel receptors. This concept of unconventional signaling is not new, but has already been described in *C. elegans* [12,37,38] and other primitive species [39–41], and also in mammals in the postnatal period [42,43]. This concept has already motivated the prediction of connection polarity instead of neuron polarity, yet on a subcircuit level [28]. By developing a gene expression-based methodology, our work is the first attempt to predict polarities of all ionotropic chemical synapses of the *C. elegans* connectome.

A surprisingly high proportion of synapses were predicted to have a complex, i.e. both excitatory and inhibitory polarity. This is due to parallel expression of cationic and anionic receptor genes – often for the same neurotransmitter – in half of the neurons. This suggests a highly complex functioning of neuronal connections that extends beyond the permanently exclusive concept of excitation-inhibition dichotomy. Since our work is based on gene expression rather than protein expression data, predictions made are derived from genetic permissibility rather than direct receptor presence. Obviously, a single synapse cannot simultaneously be excitatory and inhibitory, therefore the predicted “complexity” can be resolved mainly in two physiologically well-established ways. One is that postsynaptic receptors are not homogenously distributed across the plasma membrane but their subcellular localization is regulated. This allows the receptors to act independently [45–50] and allows the same neurotransmitter to excite and inhibit at distinct postsynaptic sites. For example, such mechanisms have been identified in the AIA [51–53] and AIB neuron groups [54]. Another explanation of “complexity” is the dynamic change of gene expression in time which is observed all through the life-span of a worm e.g. during development, learning (synaptic plasticity), and aging [55–63]. Ultimately, changes in gene expression can lead to neurotransmitter-switching and consequential up- and downregulation of receptors of opposing polarity [64,65].

Currently, there is not enough data to address either the spatial or the temporal aspects of receptor expression regulation on the worm-scale. As expression-profiling methods will provide whole-brain and dynamic proteomics data of subcellular resolution, complex synapses might be further resolved.

This paper covers neurotransmitter-mediated ionotropic connections, which account for the fast-acting system of the nervous system. Other, generally slower-acting components of the signaling system, such as G-protein coupled metabotropic or extrasynaptic signal transmission were not studied but can be targets of future research. Although the strength of prediction of our work is generally acceptable (>70%), as new data of connectivity and gene expression emerge, our method can be used to provide more accurate predictions of synaptic polarity.

Within the scope of our aims and subject to the limitations of the available connectome and gene expression data [2,21,55,66], we predicted synaptic polarities of ionotropic chemical connections in the complete *C. elegans* neuronal network, for the first time. We developed and applied a novel method that combines connectivity with presynaptic and postsynaptic gene expression data and made its tool available for users at the website http://EleganSign.linkgroup.hu/. Importantly, the natural balance of excitatory and inhibitory connections can be well approximated only if one considers both pre- and postsynaptic gene expression, a concept that was lacking from previous work. Our method opens a way to include spatial and temporal dynamics of synaptic polarity that would add a new dimension of plasticity in the excitatory:inhibitory balance. When sufficient data is available, our polarity prediction method can be applied to any neuronal (and as a concept non-neuronal) network.

## Methods

### Description of *C. elegans* connectome data

Connectome reconstruction of the adult hermaphrodite worm published by WormWiring (http://wormwiring.org) consists of 3,638 chemical connections and 2,167 gap junctions, connecting 300 neurons (the two canal-associated neurons, CANL and CANR remained isolated in this reconstruction, and therefore were omitted from the connectome-related analyses). Each of the connections have 4 attributes: the presynaptic neuron, the postsynaptic neuron, the type of the connection (chemical or electrical), and the number of synapses. The chemical connections subset consisting of 20,589 synapses connecting 297 neurons was used in our work (the sensory neuron pair PLML and PLMR, and the pharyngeal neuron M5 is isolated in the chemical synapse network). Additionally, two other connectome reconstructions – both covering a smaller number of neurons and synapses – were used for validation (Table 5 in S1 Supporting Information).

### Description of gene expression data and processing

Initially, binary gene expression data was obtained from a previous publication based on Wormbase [19]. This was extended with data of neuronal neurotransmitter [17,18] and receptor [22] expression, and with expression data from the recently published CenGen database [21] after transformation to binary information (Text 1 and Table 6 in S1 Supporting Information). For receptor expression scoring, only the genes coding ionotropic receptor subunits were evaluated according to the six functional classes based on their suggested neurotransmitter ligand (glutamate, acetylcholine or GABA) and putative ion channel type (cationic or anionic, i.e. excitatory or inhibitory). Expression of one or more genes in a functional class was considered positive.

### Prediction of synaptic polarities

Polarities were predicted for connections based on presynaptic neurotransmitter and postsynaptic receptor expression data, using nested logical and conditional formulas. In case of the method referenced as the *NT*+*R method* throughout the paper, synapses were predicted as *excitatory* or *inhibitory* if only cation channel or only anion channel receptor genes matched the presynaptic neurotransmitter, respectively; *complex* if both types of receptor genes matched; and *unpredicted* if no receptor gene matched. Alternative prediction methods were used according to the following rules. *NT-only method*: synapses were predicted *excitatory* or *inhibitory* if the presynaptic neurotransmitter was acetylcholine and/or glutamate or GABA, respectively; *complex* if acetylcholine and/or glutamate and GABA; and *unpredicted* if the neurotransmitter was none of these. *R-only method:* synapses were predicted *excitatory* or *inhibitory* if the postsynaptic receptor genes expressed were only cation channel or anion channel coding, respectively; *complex* if both types of ion channel receptor genes were expressed; and *unpredicted* if no receptor gene was expressed. Exact formulas are available in Supplementary Data.

### Software and data

Data were processed and predictions were made using Microsoft Excel (ver. 16.32) and R (RStudio 1.1.456) using standard packages.

### Data availability

Original data are available as Supplementary Information and Supplementary Data. The sign prediction tool is available at http://EleganSign.linkgroup.hu. Scripts are available on GitHub [https://github.com/bank-fenyves/CeConn-SignPrediction].

## Supporting information

S1 Data. Prediction of synaptic signs based on neurotransmitter and receptor expression data

S2 Data. Prediction of synaptic signs based on neurotransmitter expression data

S3 Data. Prediction of synaptic signs based on neurotransmitter receptor expression data

S4 Data. Prediction of synaptic and edge signs based on neurotransmitter and receptor expression data in different subnetworks

S5 Data. Prediction of synaptic signs based on neurotransmitter and receptor expression data (alternative connectome reconstructions)

## Acknowledgments

We thank members of the LINK network science group (http://linkgroup.hu) for their helpful comments. This work was supported by the Hungarian National Research, Development and Innovation Office [K131458 to P.C. and K116525 to C.S.], by the Higher Education Institutional Excellence Programme of the Ministry of Human Capacities in Hungary, within the framework of the Molecular Biology thematic programmes of the Semmelweis University, as well as by the Artificial Intelligence Research Field Excellence Programme of the National Research, Development and Innovation Office of the Ministry of Innovation and Technology in Hungary (TKP/ITM/NKFIH). B.G.F was supported by the Human Capacities Grant Management Office in Hungary [NTP-NFTÖ-18-B-0179 and NTP-NFTÖ-19-B-0264]. C.S. is a Merit Prize recipient of the Semmelweis University.

## Author contribution

B.G.F contributed to conception, design, acquisition, analysis and wrote the manuscript; G.S.S contributed to acquisition, data analysis, and to the manuscript; Z.V. contributed to the interpretation of data and provided the web-surface of the tool; C.S. contributed to interpretation of data; P.C. contributed to conception, interpretation of data, and to the manuscript.

## Competing interests

The authors declare no competing interests.

Correspondence and requests for materials should be addressed to P.C.

## Supporting Information

**S1 Supporting Information. Supplemental text and tables.**

**S1 Fig.**
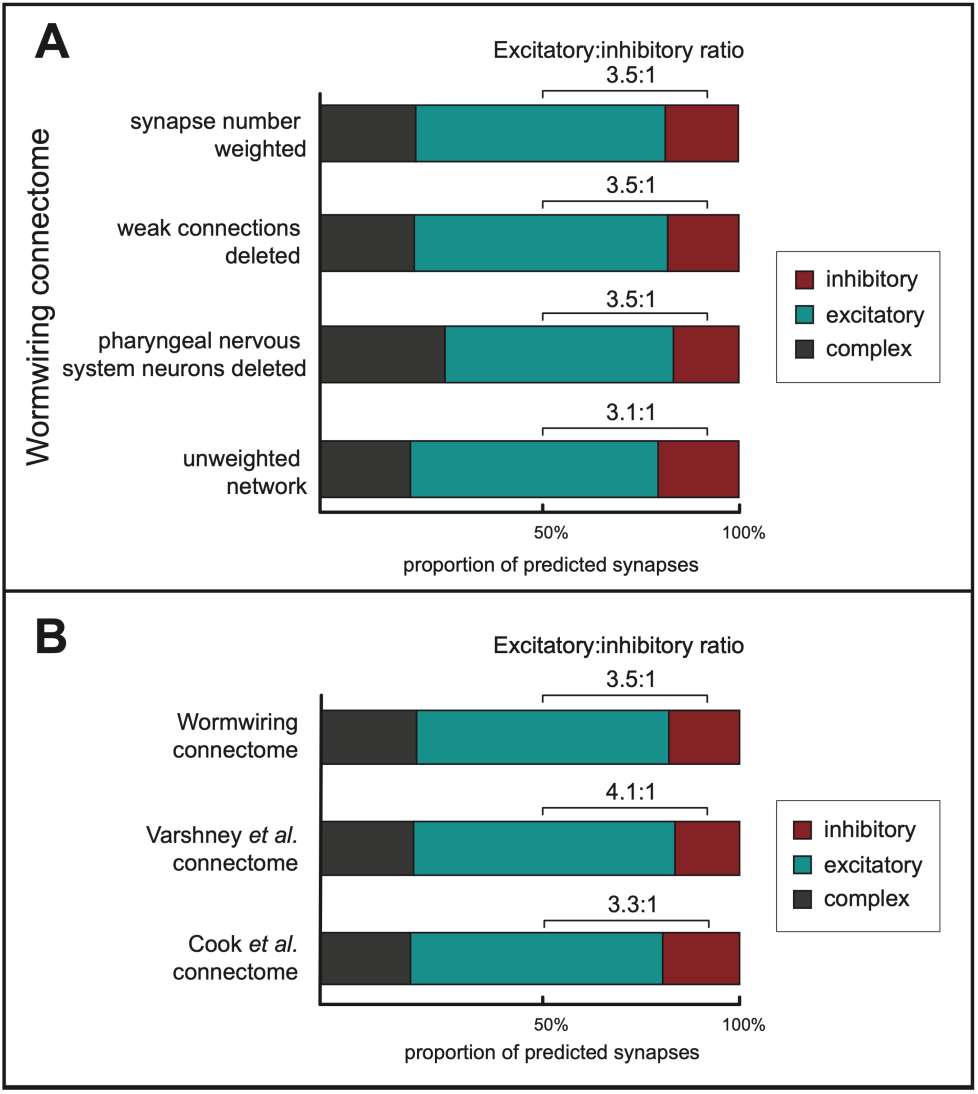
Proportions of predicted synaptic polarities in alternative *C. elegans* neuronal networks. Predictions were made based on the neurotransmitter and receptor gene expression patterns of the presynaptic and postsynaptic neurons, respectively (*NT*+*R method*, see Methods). Red, blue, and grey colors mark inhibitory, excitatory, and complex polarities, respectively. **(A)** Excitatory-inhibitory balances in alternative networks of the WormWiring connectome reconstruction. Bars from top to bottom: 1. synapse weighted network for comparative purpose (same as on Figure 3A); 2. weak links (defined by synapse number of 1) deleted [3]; 3. links connecting any of the pharyngeal nervous system neurons deleted. The rationale is that many previous work analyzed the connectome without the pharyngeal nervous system [2,67]; 4. unweighted network. **(B)** Predicted synaptic polarities for two connectome reconstructions other than Wormwiring, covering a variable number of neurons and synapses [2,3] (Table 5 in S1 Supporting Information). In summary, excitatory:inhibitory sign-balance ratios were similar in all cases, ranging between 3.1 – 4.1. Source data are provided in S1, S4 and S5 Data.

**S1 Data. Prediction of synaptic signs based on neurotransmitter and receptor expression data**

**S2 Data. Prediction of synaptic signs based on neurotransmitter expression data**

**S3 Data. Prediction of synaptic signs based on neurotransmitter receptor expression data**

**S4 Data. Prediction of synaptic and edge signs based on neurotransmitter and receptor expression data in different subnetworks**

**S5 Data. Prediction of synaptic signs based on neurotransmitter and receptor expression data (alternative connectome reconstructions)**

All above Supplementary Data files are available here: http://linkgroup.hu/elegansdata.php

## Supporting Information

### Supporting Text

#### Text 1. Gene expression

The primary sources of gene expression data used in this work [1,2] contained binary (i.e. ‘expressed’ or ‘not expressed’) information. Therefore, any extension of gene expression data needed to be carried out in a binary fashion as well. In case of the CenGEN database [3] where relative expression levels were published, data was transformed into binary information using the following rules: criteria for positive expression was avg_logFC > 0 and p_val_adj < 0.05. This way, genes that were found significantly “over-expressed” in a cell cluster compared to the rest of the cells were considered as “expressed” in that cell cluster, while the other genes were not.

Our approach of gene expression data extension was intentionally more sensitive than specific. Thus, in case of conflicting information (i.e. positive expression in one dataset and no expression data in another) evidence of positive expression was considered. Although, this method might have resulted in some false positive predictions of synaptic polarities, the balanced number of unpredicted and complex synapses observed suggested that our method was equally sensitive and specific.

### Supporting Tables

**Table 1.**
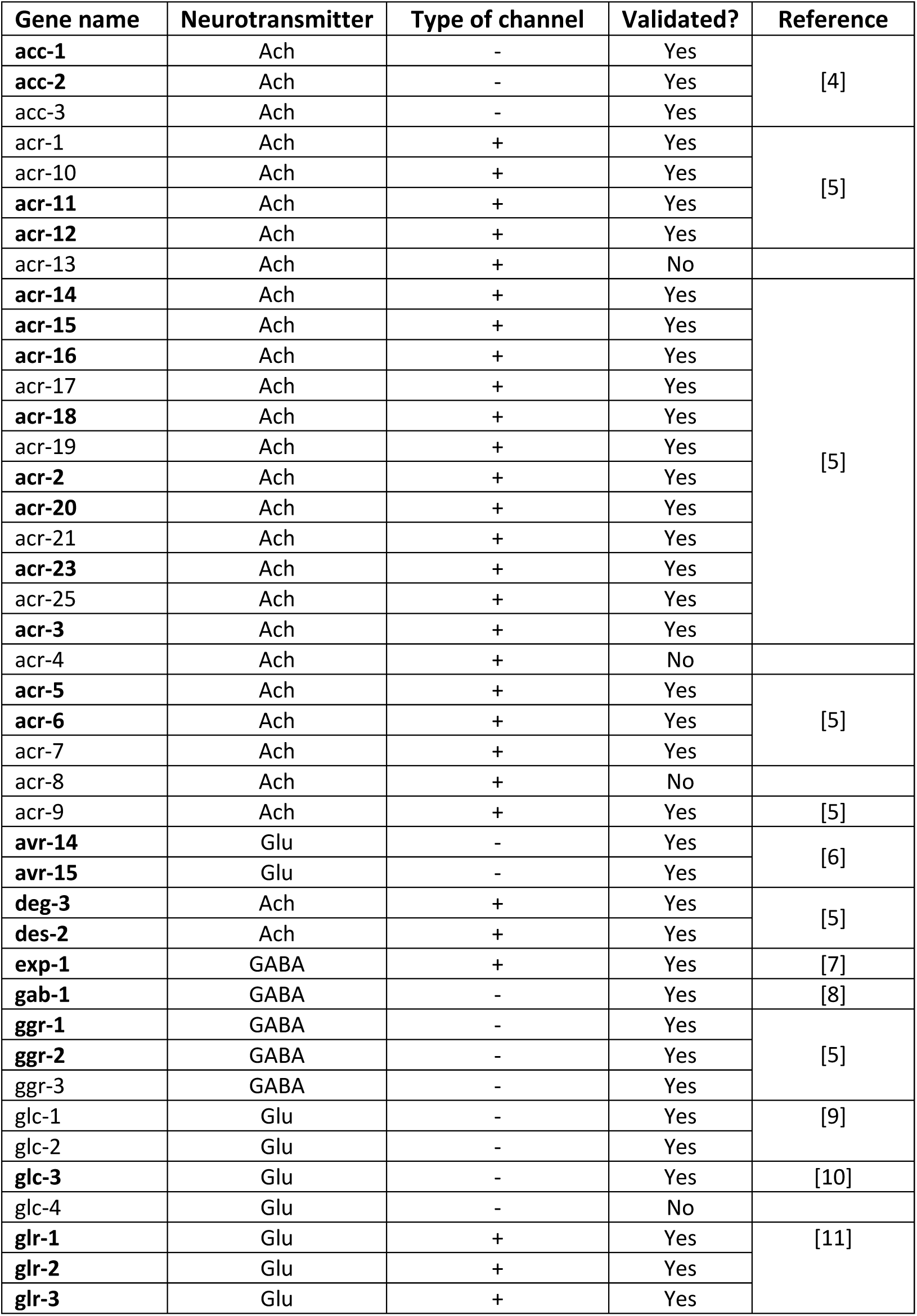
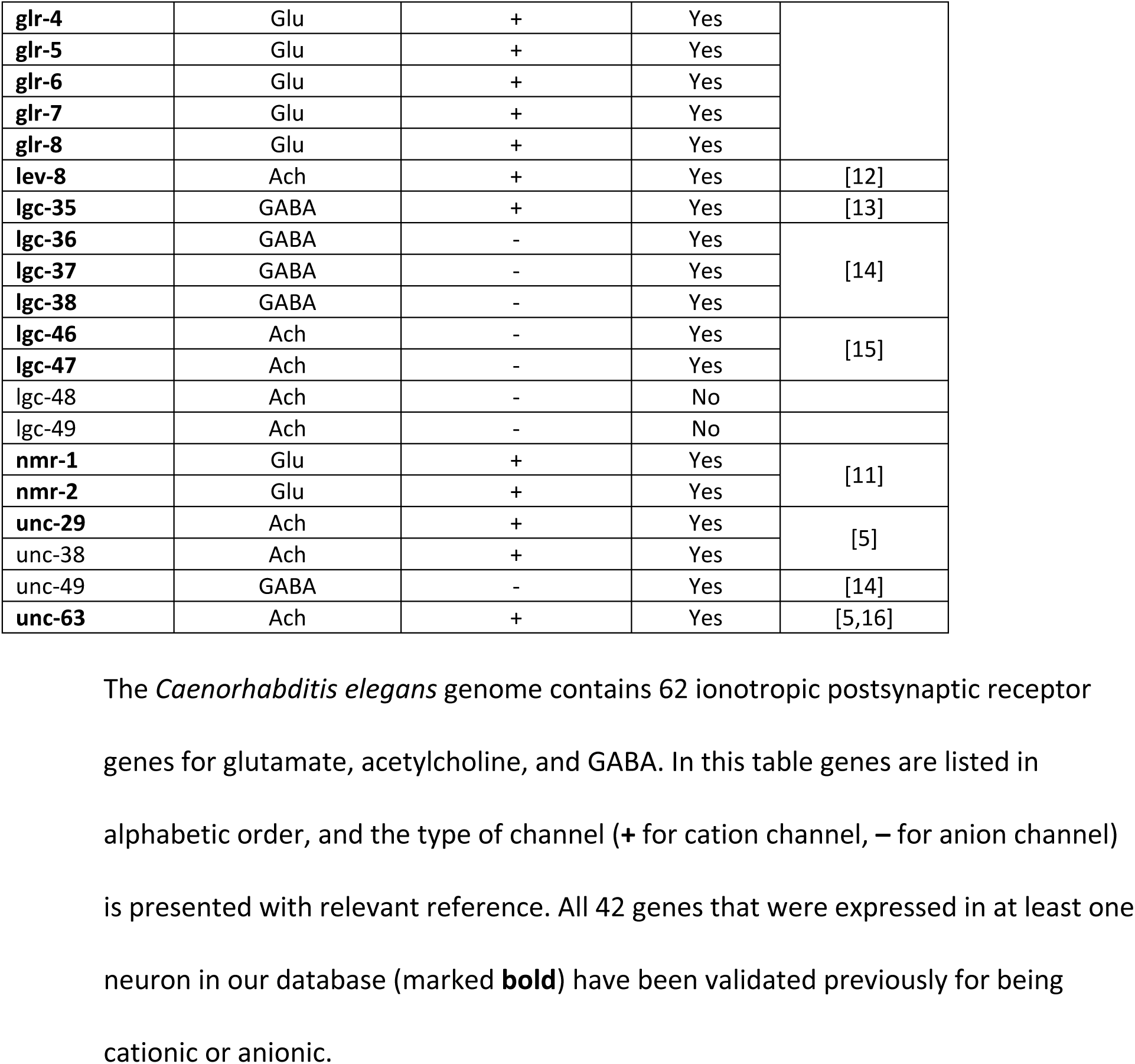
Channel type (cation or anion) of ionotropic neurotransmitter receptor genes of *C. elegans*.

**Table 2.** Distribution of neurons according to the number of ionotropic neurotransmitter receptor genes expressed.

|  |  | Number of cation channel receptor genes |  |  |  |  |  |  |  |  |  |  |
| --- | --- | --- | --- | --- | --- | --- | --- | --- | --- | --- | --- | --- |
|  |  | 0 | 1 | 2 | 3 | 4 | 5 | 6 | 7 | 8 | 9 | 10 |
| Number of anion channel receptor genes | 0 | **40** | **19** | **29** | **12** | **0** | **0** | **3** | **0** | **0** | **0** | **0** |
|  | 1 | **29** | **11** | **7** | **9** | **4** | **4** | **0** | **0** | **0** | **0** | **0** |
|  | 2 | **16** | **2** | **6** | **22** | **13** | **3** | **1** | **0** | **2** | **0** | **2** |
|  | 3 | **1** | **3** | **8** | **6** | **2** | **0** | **1** | **9** | **0** | **0** | **0** |
|  | 4 | **2** | **0** | **11** | **3** | **2** | **0** | **11** | **0** | **0** | **0** | **0** |
|  | 5 | **0** | **4** | **1** | **0** | **0** | **0** | **0** | **0** | **0** | **0** | **0** |
|  | 6 | **0** | **0** | **0** | **0** | **0** | **2** | **0** | **0** | **0** | **0** | **0** |
|  | 7 | **0** | **0** | **0** | **0** | **0** | **2** | **0** | **0** | **0** | **0** | **0** |

Neuronal gene expression database was compiled from available datasets and manually curated (Methods). Genes encoding ionotropic receptors for glutamate, acetylcholine or GABA were grouped according to the type of ion channel (cation or anion), i.e. whether being excitatory or inhibitory. Bold numbers represent number of neurons expressing certain numbers of excitatory and/or inhibitory receptor genes. 63 neurons express only cation-channel receptor genes (green), while 48 neurons express only anion-channel receptor genes (red). 151 neurons express a mixture of cation- and anion-channel receptor genes (grey). Source data is in S1 Data.

**Table 3.**
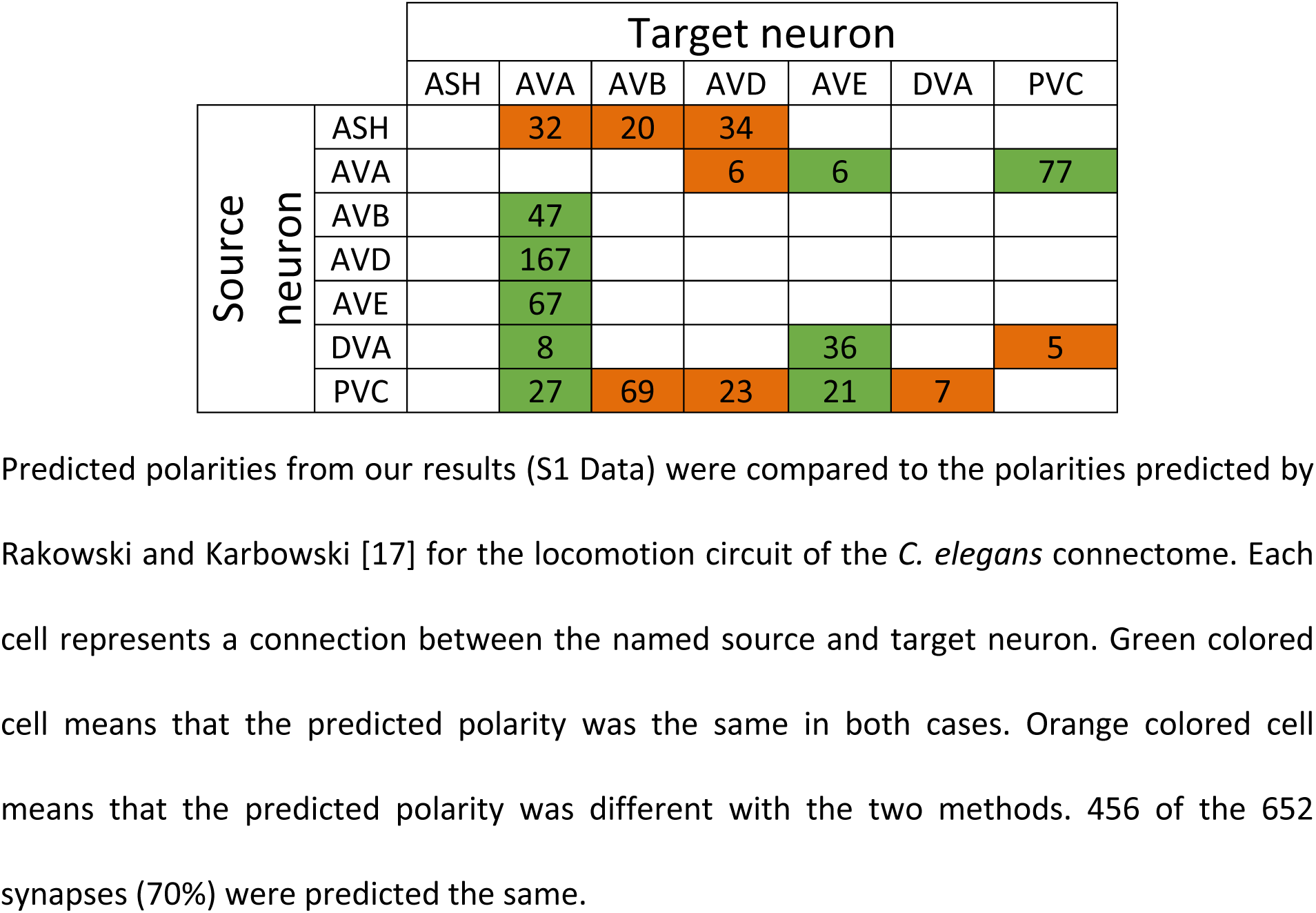
Validation of results with a previous synaptic polarity prediction paper.

|  |  | Target neuron |  |  |  |  |  |  |
| --- | --- | --- | --- | --- | --- | --- | --- | --- |
|  |  | ASH | AVA | AVB | AVD | AVE | DVA | PVC |
| Source neuron | ASH |  | 32 | 20 | 34 |  |  |  |
|  | AVA |  |  |  | 6 | 6 |  | 77 |
|  | AVB |  | 47 |  |  |  |  |  |
|  | AVD |  | 167 |  |  |  |  |  |
|  | AVE |  | 67 |  |  |  |  |  |
|  | DVA |  | 8 |  |  | 36 |  | 5 |
|  | PVC |  | 27 | 69 | 23 | 21 | 7 |  |

Predicted polarities from our results (S1 Data) were compared to the polarities predicted by Rakowski and Karbowski [17] for the locomotion circuit of the *C. elegans* connectome. Each cell represents a connection between the named source and target neuron. Green colored cell means that the predicted polarity was the same in both cases. Orange colored cell means that the predicted polarity was different with the two methods. 456 of the 652 synapses (70%) were predicted the same.

**Table 4.**
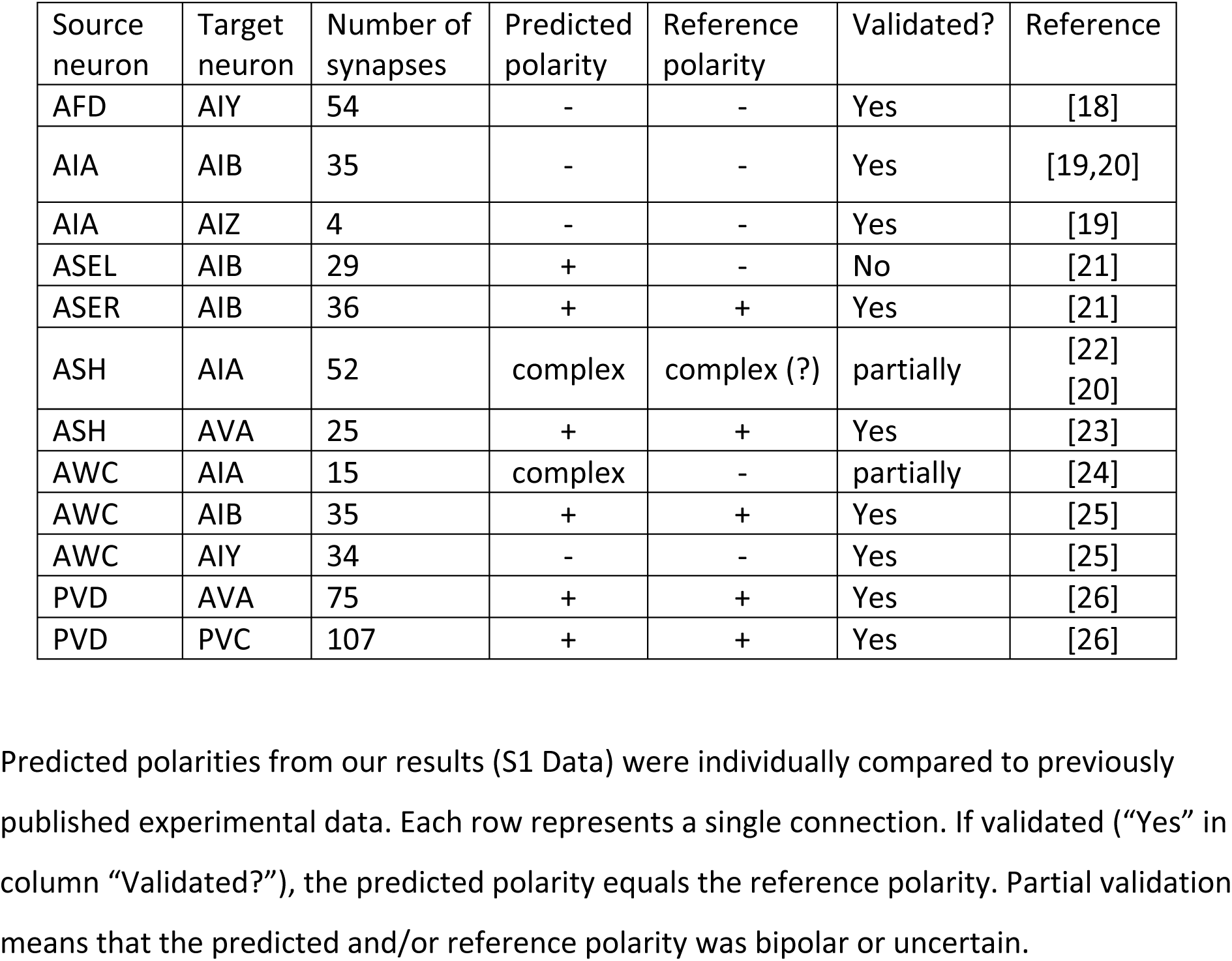
Validation of predictions with previously published experimental results.

**Table 5.**
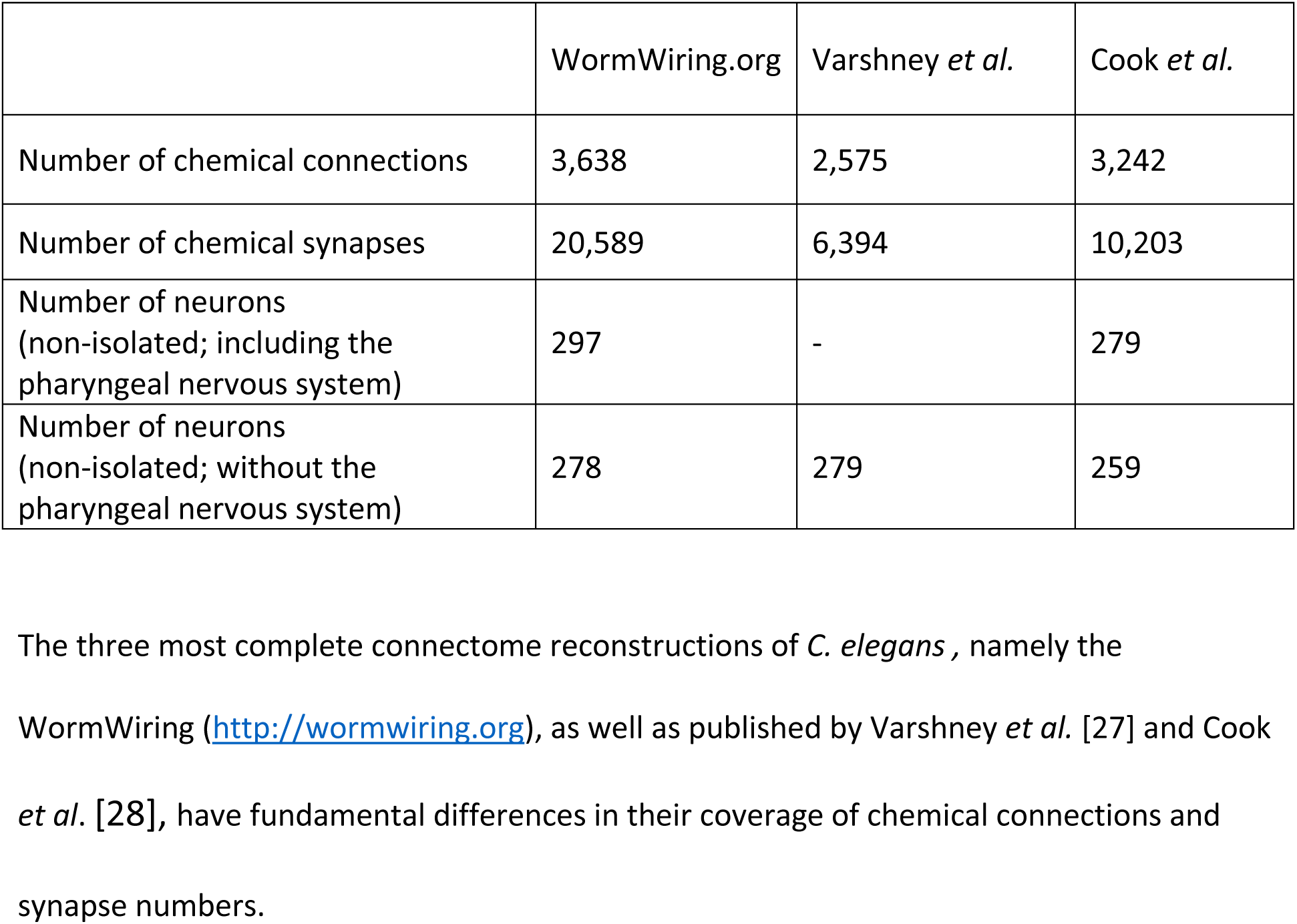
Comparison of three chemical synapse connectome reconstructions.

|  | WormWiring.org | Varshney *et al.* | Cook *et al.* |
| --- | --- | --- | --- |
| Number of chemical connections | 3,638 | 2,575 | 3,242 |
| Number of chemical synapses | 20,589 | 6,394 | 10,203 |
| Number of neurons<br>(non-isolated; including the<br>pharyngeal nervous system) | 297 | - | 279 |
| Number of neurons<br>(non-isolated; without the<br>pharyngeal nervous system) | 278 | 279 | 259 |

The three most complete connectome reconstructions of *C. elegans*, namely the WormWiring (http://wormwiring.org), as well as published by Varshney *et al.* [27] and Cook *et al.* [28], have fundamental differences in their coverage of chemical connections and synapse numbers.

**Table 6.**
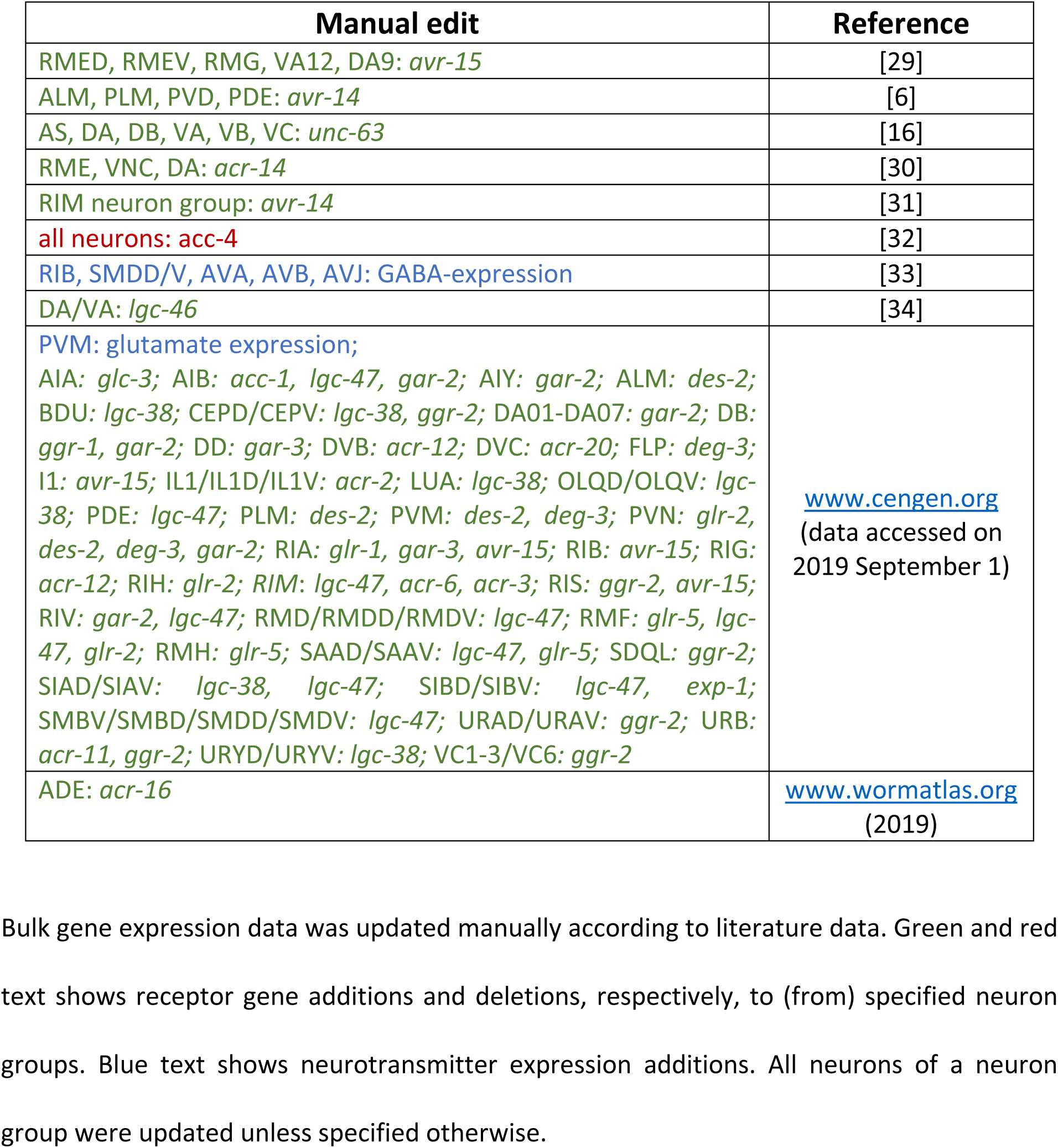
Manual curation and edits.

## References

1. White JG, Southgate E, Thomson JN, Brenner S. The structure of the nervous system of the nematode *Caenorhabditis elegans*. Philos Trans R Soc London B, Biol Sci. 1986;314: 1–340. doi: 10.1098/rstb.1986.0056

2. Varshney LR, Chen BL, Paniagua E, Hall DH, Chklovskii DB. Structural properties of the *Caenorhabditis elegans* neuronal network. PLoS Comput Biol. 2011;7: e1001066. doi: 10.1371/journal.pcbi.1001066

3. Cook SJ, Jarrell TA, Brittin CA, Wang Y, Bloniarz AE, Yakovlev MA, et al. Whole-animal connectomes of both *Caenorhabditis elegans* sexes. Nature. 2019;571: 63–71. doi: 10.1038/s41586-019-1352-7

4. Wicks SR, Roehrig CJ, Rankin CH. A dynamic network simulation of the nematode tap withdrawal circuit: predictions concerning synaptic function using behavioral criteria. J Neurosci. 1996;16: 4017–4031.

5. Rakowski F, Srinivasan J, Sternberg PW, Karbowski J. Synaptic polarity of the interneuron circuit controlling *C. elegans* locomotion. Front Comput Neurosci. 2013;7: 128. doi: 10.3389/fncom.2013.00128

6. Dong C-Y, Cho K-H. An optimally evolved connective ratio of neural networks that maximizes the occurrence of synchronized bursting behavior. BMC Syst Biol. 2012;6: 23. doi: 10.1186/1752-0509-6-23

7. McIntire SL, Jorgensen E, Kaplan J, Horvitz HR. The GABAergic nervous system of *Caenorhabditis elegans*. Nature. 1993;364: 337–341. doi: 10.1038/364337a0

8. Gendrel M, Atlas EG, Hobert O. A cellular and regulatory map of the GABAergic nervous system of *C. elegans*. Elife. 2016; e17686. doi: 10.7554/eLife.17686

9. Rubin R, Abbott LF, Sompolinsky H. Balanced excitation and inhibition are required for high-capacity, noise-robust neuronal selectivity. Proc Natl Acad Sci. 2017;114: E9366–E9375. doi: 10.1073/pnas.1705841114

10. Bhatia A, Moza S, Bhalla US. Precise excitation-inhibition balance controls gain and timing in the hippocampus. Elife. 2019;8: e43415. doi: 10.7554/eLife.43415

11. Baker C, Ebsch C, Lampl I, Rosenbaum R. Correlated states in balanced neuronal networks. Phys Rev E. 2019;99: 52414. doi: 10.1103/PhysRevE.99.052414

12. Putrenko I, Zakikhani M, Dent JA. A family of acetylcholine-gated chloride channel subunits in *Caenorhabditis elegans*. J Biol Chem. 2005;280: 6392–6398. doi: 10.1074/jbc.M412644200

13. Li Z, Liu J, Zheng M, Xu XZS. Encoding of both analog- and digital-like behavioral outputs by one *C. elegans* interneuron. Cell. 2014;159: 751–765. doi: 10.1016/j.cell.2014.09.056

14. Chalasani SH, Chronis N, Tsunozaki M, Gray JM, Ramot D, Goodman MB, et al. Dissecting a circuit for olfactory behaviour in *Caenorhabditis elegans*. Nature. 2007;450: 63–70. doi: 10.1038/nature06292

15. Marom S, Shahaf G. Development, learning and memory in large random networks of cortical neurons: lessons beyond anatomy. Q Rev Biophys. 2002;35: 63–87. doi: 10.1017/S0033583501003742

16. Pastore VP, Massobrio P, Godjoski A, Martinoia S. Identification of excitatory-inhibitory links and network topology in large-scale neuronal assemblies from multi-electrode recordings. PLOS Comput Biol. 2018;14: e1006381.

17. Loer CM, Rand JB. The evidence for classical neurotransmitters in Caenorhabditis elegans. Altun ZF, Herndon LA, editors. WormAtlas. 2016. doi: 10.3908/wormatlas.5.200

18. Pereira L, Kratsios P, Serrano-Saiz E, Sheftel H, Mayo AE, Hall DH, et al. A cellular and regulatory map of the cholinergic nervous system of *C. elegans*. Elife. 2015;4: e12432. doi: 10.7554/eLife.12432

19. Hobert O, Glenwinkel L, White J. Revisiting neuronal cell type classification in *Caenorhabditis elegans*. Curr Biol. 2016;26: R1197–R1203. doi: 10.1016/j.cub.2016.10.027

20. Serrano-Saiz E, Poole RJ, Felton T, Zhang F, De La Cruz ED, Hobert O. Modular control of glutamatergic neuronal identity in *C. elegans* by distinct homeodomain proteins. Cell. 2013;155: 659–673. doi: 10.1016/j.cell.2013.09.052

21. Taylor SR, Santpere G, Reilly M, Glenwinkel L, Poff A, McWhirter R, et al. Expression profiling of the mature *C. elegans* nervous system by single-cell RNA-sequencing. bioRxiv. 2019; 737577. doi: 10.1101/737577

22. Altun ZF. Neurotransmitter receptors in *Caenorhabditis elegans*. WormAtlas. 2011. doi: 10.3908/wormatlas.5.202

23. Martikainen MH, Kaneko KI, Hari R. Suppressed responses to self-triggered sounds in the human auditory cortex. Cereb Cortex. 2005;15: 299–302. doi: 10.1093/cercor/bhh131

24. Dalenoort G, de Vries PH. The essential role of binding for cognition in living systems. In: Schaub H, Detje F, Bruggemann U, editors. Logic af artificial life: Abstracting and synthesizing the principles of living systems. Berlin: Aka GmbH; 2004. pp. 32–39.

25. Ágg B, Császár A, Szalay-Bekő M, Veres D V., Mizsei R, Ferdinandy P, et al. The EntOptLayout Cytoscape plug-in for the efficient visualization of major protein complexes in protein–protein interaction and signalling networks. Bioinformatics. 2019;35: 4490–4492. doi: 10.1093/bioinformatics/btz257

26. Buzsáki G, Kaila K, Raichle M. Inhibition and brain work. Neuron. 2007;56: 771–783. doi: 10.1016/j.neuron.2007.11.008

27. Markram H, Toledo-Rodriguez M, Wang Y, Gupta A, Silberberg G, Wu C. Interneurons of the neocortical inhibitory system. Nat Rev Neurosci. 2004;5: 793–807. doi: 10.1038/nrn1519

28. Rakowski F, Karbowski J. Optimal synaptic signaling connectome for locomotory behavior in *Caenorhabditis elegans*: Design minimizing energy cost. PLOS Comput Biol. 2017;13: e1005834.

29. Sohal VS, Rubenstein JLR. Excitation-inhibition balance as a framework for investigating mechanisms in neuropsychiatric disorders. Mol Psychiatry. 2019/05/14. 2019;24: 1248–1257. doi: 10.1038/s41380-019-0426-0

30. Blaszczyk JW. Parkinson’s Disease and neurodegeneration: GABA-collapse hypothesis. Front Neurosci. 2016;10: 269. doi: 10.3389/fnins.2016.00269

31. Liu G. Local structural balance and functional interaction of excitatory and inhibitory synapses in hippocampal dendrites. Nat Neurosci. 2004;7: 373–379. doi: 10.1038/nn1206

32. Markram H, Muller E, Ramaswamy S, Reimann MW, Abdellah M, Sanchez CA, et al. Reconstruction and simulation of neocortical microcircuitry. Cell. 2015;163: 456–492. doi: 10.1016/j.cell.2015.09.029

33. Gulyás AI, Megías M, Emri Z, Freund TF. Total number and ratio of excitatory and inhibitory synapses converging onto single interneurons of different types in the CA1 area of the rat hippocampus. J Neurosci. 1999;19: 10082–10097. doi: 10.1523/JNEUROSCI.19-22-10082.1999

34. Nakanishi K, Kukita F. Intracellular [Cl-] modulates synchronous electrical activity in rat neocortical neurons in culture by way of GABAergic inputs. Brain Res. 2000;863: 192–204. doi: 10.1016/S0006-8993(00)02152-1

35. Leskovec J, Huttenlocher D, Kleinberg J. Signed networks in social media. Proceedings of the SIGCHI Conference on Human Factors in Computing Systems. New York, NY, USA: ACM; 2010. pp. 1361–1370. doi: 10.1145/1753326.1753532

36. Kirkley A, Cantwell GT, Newman MEJ. Balance in signed networks. Phys Rev E. 2019;99: 012320. doi: 10.1103/PhysRevE.99.012320

37. Beg AA, Jorgensen EM. EXP-1 is an excitatory GABA-gated cation channel. Nat Neurosci. 2003;6: 1145–1152. doi: 10.1038/nn1136

38. Martinez-Torres A, Miledi R. Expression of *Caenorhabditis elegans* neurotransmitter receptors and ion channels in Xenopus oocytes. Proc Natl Acad Sci. 2006;103: 5120–5124. doi: 10.1073/pnas.0600739103

39. Kehoe J, McIntosh JM. Two distinct nicotinic receptors, one pharmacologically similar to the vertebrate α7-containing receptor, mediate Cl currents in *Aplysia* neurons. J Neurosci. 1998;18: 8198–8213. doi: 10.1523/JNEUROSCI.18-20-08198.1998

40. Cully DF, Paress PS, Liu KK, Schaeffer JM, Arena JP. Identification of a *Drosophila melanogaster* glutamate-gated chloride channel sensitive to the antiparasitic agent avermectin. J Biol Chem. 1996;271: 20187–20191. doi: 10.1074/jbc.271.33.20187

41. Liu WW, Wilson RI. Glutamate is an inhibitory neurotransmitter in the *Drosophila* olfactory system. Proc Natl Acad Sci U S A. 2013;110: 10294–10299. doi: 10.1073/pnas.1220560110

42. Kullmann PHM, Ene FA, Kandler K. Glycinergic and GABAergic calcium responses in the developing lateral superior olive. Eur J Neurosci. 2002;15: 1093–1104. doi: 10.1046/j.1460-9568.2002.01946.x

43. Wolstenholme AJ. Glutamate-gated chloride channels. J Biol Chem. 2012;287: 40232–40238. doi: 10.1074/jbc.R112.406280

44. Zhou D, Rangan A V, McLaughlin DW, Cai D. Spatiotemporal dynamics of neuronal population response in the primary visual cortex. Proc Natl Acad Sci. 2013;110: 9517–9522. doi: 10.1073/pnas.1308167110

45. Tao L, Porto D, Li Z, Fechner S, Lee SA, Goodman MB, et al. Parallel processing of two mechanosensory modalities by a single neuron in *C. elegans*. Dev Cell. 2019;51: 543–658. doi: 10.1016/j.devcel.2019.10.008

46. Nusser Z. Subcellular distribution of neurotransmitter receptors and voltage-gated ion channels. In: Stuart G, Spruston N, Hausser M, editors. Dendrites. Oxford University Press; 2012. pp. 154–188. doi: 10.1093/acprof:oso/9780198566564.003.0007

47. Megías M, Emri Z, Freund TF, Gulyás AI. Total number and distribution of inhibitory and excitatory synapses on hippocampal CA1 pyramidal cells. Neuroscience. 2001;102: 527–540. doi: 10.1016/S0306-4522(00)00496-6

48. Zou W, Fu J, Zhang H, Du K, Huang W, Yu J, et al. Decoding the intensity of sensory input by two glutamate receptors in one *C. elegans* interneuron. Nat Commun. 2018;9: 4311. doi: 10.1038/s41467-018-06819-5

49. Arey RN, Kaletsky R, Murphy CT. Nervous system-wide profiling of presynaptic mRNAs reveals regulators of associative memory. Sci Rep. 2019;9: 20314. doi: 10.1038/s41598-019-56908-8

50. Stetak A, Hörndli F, Maricq A V, van den Heuvel S, Hajnal A. Neuron-specific regulation of associative learning and memory by MAGI-1 in *C. elegans*. PLoS One. 2009;4: e6019. doi: 10.1371/journal.pone.0006019

51. Choi S, Taylor KP, Chatzigeorgiou M, Hu Z, Schafer WR, Kaplan JM. Sensory neurons arouse *C. elegans* locomotion via both glutamate and neuropeptide release. PLOS Genet. 2015;11: e1005359. doi: 10.1371/journal.pgen.1005359

52. Shinkai Y, Yamamoto Y, Fujiwara M, Tabata T, Murayama T, Hirotsu T, et al. Behavioral choice between conflicting alternatives is regulated by a receptor guanylyl cyclase, GCY-28, and a receptor tyrosine kinase, SCD-2, in AIA interneurons of *Caenorhabditis elegans*. J Neurosci. 2011;31: 3007–3015. doi: 10.1523/JNEUROSCI.4691-10.2011

53. Chalasani SH, Kato S, Albrecht DR, Nakagawa T, Abbott LF, Bargmann CI. Neuropeptide feedback modifies odor-evoked dynamics in *Caenorhabditis elegans* olfactory neurons. Nat Neurosci. 2010;13: 615–621. doi: 10.1038/nn.2526

54. Kuramochi M, Doi M. An excitatory/inhibitory switch from asymmetric sensory neurons defines postsynaptic tuning for a rapid response to NaCl in *Caenorhabditis elegans*. Front Mol Neurosci. 2019;11: 484. doi: 10.3389/fnmol.2018.00484

55. Witvliet D, Mulcahy B, Mitchell JK, Meirovitch Y, Berger DK, Wu Y, et al. Connectomes across development reveal principles of brain maturation in *C. elegans*. bioRxiv. 2020. doi: 10.1101/2020.04.30.066209

56. Serrano-Saiz E, Pereira L, Gendrel M, Aghayeva U, Battacharya A, Howell K, et al. A neurotransmitter atlas of the *Caenorhabditis elegans* male nervous system reveals sexually dimorphic neurotransmitter usage. Genetics. 2017;206: 1251–1269. doi: 10.1534/genetics.117.202127

57. Kunitomo H, Sato H, Iwata R, Satoh Y, Ohno H, Yamada K, et al. Concentration memory-dependent synaptic plasticity of a taste circuit regulates salt concentration chemotaxis in *Caenorhabditis elegans*. Nat Commun. 2013;4: 2210. doi: 10.1038/ncomms3210

58. Ho VM, Lee J-A, Martin KC. The cell biology of synaptic plasticity. Science. 2011;334: 623–628. doi: 10.1126/science.1209236

59. Hadziselimovic N, Vukojevic V, Peter F, Milnik A, Fastenrath M, Fenyves BG, et al. Forgetting is regulated via Musashi-mediated translational control of the Arp2/3 complex. Cell. 2014;156: 1153–1166. doi: 10.1016/j.cell.2014.01.054

60. Ingrosso A, Abbott LF. Training dynamically balanced excitatory-inhibitory networks. PLoS One. 2019;14: e0220547. doi: 10.1371/journal.pone.0220547

61. Freytag V, Probst S, Hadziselimovic N, Boglari C, Hauser Y, Peter F, et al. Genome-wide temporal expression profiling in *Caenorhabditis elegans* identifies a core gene set related to long-term memory. J Neurosci. 2017;37: 6661–6672. doi: 10.1523/JNEUROSCI.3298-16.2017

62. Hangya B, Ranade SP, Lorenc M, Kepecs A. Central cholinergic neurons are rapidly recruited by reinforcement feedback. Cell. 2015;162: 1155–1168. doi: 10.1016/j.cell.2015.07.057

63. Hoerndli FJ, Walser M, Fröhli Hoier E, de Quervain D, Papassotiropoulos A, Hajnal A. A conserved function of *C. elegans* CASY-1 calsyntenin in associative learning. PLoS One. 2009;4: e4880. doi: 10.1371/journal.pone.0004880

64. Hammond-Weinberger DR, Wang Y, Glavis-Bloom A, Spitzer NC. Mechanism for neurotransmitter-receptor matching. Proc Natl Acad Sci. 2020;117: 4368–4374. doi: 10.1073/pnas.1916600117

65. Spitzer NC. Neurotransmitter switching in the developing and adult brain. Annu Rev Neurosci. 2017;40: 1–19. doi: 10.1146/annurev-neuro-072116-031204

66. Yemini E, Lin A, Nejatbakhsh A, Varol E, Sun R, Mena GE, et al. NeuroPAL: A neuronal polychromatic atlas of landmarks for whole-brain imaging in *C. elegans*. bioRxiv. 2019; 676312. doi: 10.1101/676312

67. Milo R, Shen-Orr S, Itzkovitz S, Kashtan N, Chklovskii D, Alon U. Network motifs: simple building blocks of complex networks. Science. 2002;298: 824–827. doi: 10.1126/science.298.5594.824

## Supplementary References

1. Hobert O. A map of terminal regulators of neuronal identity in *Caenorhabditis elegans*. Wiley Interdisciplinary Reviews: Developmental Biology. John Wiley & Sons, Inc.; 2016. pp. 474–498. doi: 10.1002/wdev.233

2. Altun ZF. Neurotransmitter receptors in *Caenorhabditis elegans*. WormAtlas. 2011. doi: 10.3908/wormatlas.5.202

3. Taylor SR, Santpere G, Reilly M, Glenwinkel L, Poff A, McWhirter R, et al. Expression profiling of the mature *C. elegans* nervous system by single-cell RNA-sequencing. bioRxiv. 2019; 737577. doi: 10.1101/737577

4. Putrenko I, Zakikhani M, Dent JA. A family of acetylcholine-gated chloride channel subunits in *Caenorhabditis elegans*. J Biol Chem. 2005;280: 6392–6398. doi: 10.1074/jbc.M412644200

5. Jones A, Sattelle D. The cys-loop ligand-gated ion channel gene superfamily of the nematode, *Caenorhabditis elegans*. Invert Neurosci. 2008;8: 41–47. doi: 10.1007/s10158-008-0068-4

6. Dent JA, Smith MM, Vassilatis DK, Avery L. The genetics of ivermectin resistance in *Caenorhabditis elegans*. Proc Natl Acad Sci. 2000;97: 26742679. doi: 10.1073/pnas.97.6.2674

7. Beg AA, Jorgensen EM. EXP-1 is an excitatory GABA-gated cation channel. Nat Neurosci. 2003;6: 1145–1152. doi: 10.1038/nn1136

8. Feng X-P, Hayashi J, Beech R, Prichard R. Study of the nematode putative GABA type-A receptor subunits: Evidence for modulation by ivermectin. J Neurochem. 2002;83: 870–878. doi: 10.1046/j.1471-4159.2002.01199.x

9. Cully DF, Vassilatis DK, Liu KK, Paress PS, Van der Ploeg LHT, Schaeffer JM, et al. Cloning of an avermectin-sensitive glutamate-gated chloride channel from *Caenorhabditis elegans*. Nature. 1994;371: 707–711. doi: 10.1038/371707a0

10. Horoszok L, Raymond V, Sattelle DB, Wolstenholme AJ. GLC-3: a novel fipronil and BIDN-sensitive, but picrotoxinin-insensitive, L-glutamate-gated chloride channel subunit from *Caenorhabditis elegans*. Br J Pharmacol. 2001;132: 1247–1254. doi: 10.1038/sj.bjp.0703937

11. Brockie PJ, Madsen DM, Zheng Y, Mellem J, Maricq A V. Differential expression of glutamate receptor subunits in the nervous system of *Caenorhabditis elegans* and their regulation by the homeodomain protein UNC-42. J Neurosci. 2001;21: 1510–1522. doi: 10.1523/JNEUROSCI.21-05-01510.2001

12. Towers P, Edwards B, Richmond J, Sattelle D. The *Caenorhabditis elegans* lev-8 gene encodes a novel type of nicotinic acetylcholine receptor α subunit. J Neurochem. 2005;93: 1–9. doi: 10.1111/j.1471-4159.2004.02951.x

13. Nicholl GCB, Jawad AK, Weymouth R, Zhang H, Beg AA. Pharmacological characterization of the excitatory ‘Cys-loop’ GABA receptor family in *Caenorhabditis elegans*. Br J Pharmacol. 2017;174: 781–795. doi: 10.1111/bph.13736

14. Bamber BA, Beg AA, Twyman RE, Jorgensen EM. The *Caenorhabditis elegans* unc-49 locus encodes multiple subunits of a heteromultimeric GABA receptor. J Neurosci. 1999;19: 5348–5359. doi: 10.1523/JNEUROSCI.19-13-05348.1999

15. Hobert O, Glenwinkel L, White J. Revisiting neuronal cell type classification in *Caenorhabditis elegans*. Curr Biol. 2016;26: R1197–R1203. doi: 10.1016/j.cub.2016.10.027

16. Culetto E, Baylis HA, Richmond JE, Jones AK, Fleming JT, Squire MD, et al. The *Caenorhabditis elegans* unc-63 gene encodes a levamisole-sensitive nicotinic acetylcholine receptor α subunit. J Biol Chem. 2004;279: 42476–42483. doi: 10.1074/jbc.M404370200

17. Rakowski F, Karbowski J. Optimal synaptic signaling connectome for locomotory behavior in *Caenorhabditis elegans*: Design minimizing energy cost. PLOS Comput Biol. 2017;13: e1005834.

18. Narayan A, Laurent G, Sternberg PW. Transfer characteristics of a thermosensory synapse in *Caenorhabditis elegans*. Proc Natl Acad Sci. 2011;108: 9667–9672. doi: 10.1073/pnas.1106617108

19. Wakabayashi T, Kitagawa I, Shingai R. Neurons regulating the duration of forward locomotion in *Caenorhabditis elegans*. Neurosci Res. 2004;50: 103–111. doi: 10.1016/j.neures.2004.06.005

20. Shinkai Y, Yamamoto Y, Fujiwara M, Tabata T, Murayama T, Hirotsu T, et al. Behavioral choice between conflicting alternatives is regulated by a receptor guanylyl cyclase, GCY-28, and a receptor tyrosine kinase, SCD-2, in AIA interneurons of *Caenorhabditis elegans*. J Neurosci. 2011;31: 3007–3015. doi: 10.1523/JNEUROSCI.4691-10.2011

21. Kuramochi M, Doi M. An excitatory/inhibitory switch from asymmetric sensory neurons defines postsynaptic tuning for a rapid response to NaCl in *Caenorhabditis elegans*. Front Mol Neurosci. 2019;11: 484. doi: 10.3389/fnmol.2018.00484

22. Choi S, Taylor KP, Chatzigeorgiou M, Hu Z, Schafer WR, Kaplan JM. Sensory neurons arouse *C. elegans* locomotion via both glutamate and neuropeptide release. PLOS Genet. 2015;11: e1005359. doi: 10.1371/journal.pgen.1005359

23. Lindsay TH, Thiele TR, Lockery SR. Optogenetic analysis of synaptic transmission in the central nervous system of the nematode *Caenorhabditis elegans*. Nat Commun. 2011;2: 306–309. doi: 10.1038/ncomms1304

24. Chalasani SH, Kato S, Albrecht DR, Nakagawa T, Abbott LF, Bargmann CI. Neuropeptide feedback modifies odor-evoked dynamics in *Caenorhabditis elegans* olfactory neurons. Nat Neurosci. 2010;13: 615–621. doi: 10.1038/nn.2526

25. Chalasani SH, Chronis N, Tsunozaki M, Gray JM, Ramot D, Goodman MB, et al. Dissecting a circuit for olfactory behaviour in *Caenorhabditis elegans*. Nature. 2007;450: 63–70. doi: 10.1038/nature06292

26. Husson SJ, Gottschalk A, Leifer AM. Optogenetic manipulation of neural activity in *C. elegans*: From synapse to circuits and behaviour. Biol Cell. 2013;105: 235–250. doi: 10.1111/boc.201200069

27. Varshney LR, Chen BL, Paniagua E, Hall DH, Chklovskii DB. Structural properties of the *Caenorhabditis elegans* neuronal network. PLoS Comput Biol. 2011;7: e1001066. doi: 10.1371/journal.pcbi.1001066

28. Cook SJ, Jarrell TA, Brittin CA, Wang Y, Bloniarz AE, Yakovlev MA, et al. Whole-animal connectomes of both *Caenorhabditis elegans* sexes. Nature. 2019;571: 63–71. doi: 10.1038/s41586-019-1352-7

29. Dent JA, Davis MW, Avery L. *avr-15* encodes a chloride channel subunit that mediates inhibitory glutamatergic neurotransmission and ivermectin sensitivity in *Caenorhabditis elegans*. EMBO J. 1997;16: 5867–5879. doi: 10.1093/emboj/16.19.5867

30. Fox RM, Von Stetina SE, Barlow SJ, Shaffer C, Olszewski KL, Moore JH, et al. A gene expression fingerprint of *C. elegans* embryonic motor neurons. BMC Genomics. 2005;6: 42. doi: 10.1186/1471-2164-6-42

31. Piggott BJ, Liu J, Feng Z, Wescott SA, Xu XZS. The neural circuits and synaptic mechanismsm underlying motor initiation in *C. elegans*. Cell. 2011;147: 922–933. doi: 10.1016/j.cell.2011.08.053

32. Pereira L, Kratsios P, Serrano-Saiz E, Sheftel H, Mayo AE, Hall DH, et al. A cellular and regulatory map of the cholinergic nervous system of *C. elegans*. Elife. 2015;4: e12432. doi: 10.7554/eLife.12432

33. Gendrel M, Atlas EG, Hobert O. A cellular and regulatory map of the GABAergic nervous system of *C. elegans*. Elife. 2016; e17686. doi: 10.7554/eLife.17686

34. Liu P, Chen B, Mailler R, Wang Z-W. Antidromic-rectifying gap junctions amplify chemical transmission at functionally mixed electrical-chemical synapses. Nat Commun. 2017;8: 14818. doi: 10.1038/ncomms14818

